# Cells from the same lineage switch from reduction to enhancement of size variability in *Arabidopsis* sepals

**DOI:** 10.1101/132431

**Authors:** Satoru Tsugawa, Nathan Hervieux, Daniel Kierzkowski, Anne-Lise Routier-Kierzkoswska, Aleksandra Sapala, Olivier Hamant, Richard S. Smith, Adrienne H. K. Roeder, Arezki Boudaoud, Chun-Biu Li

## Abstract

Organs form with remarkably consistent sizes and shapes during development, whereas a high variability in size and growth is observed at cell level. Given this contrast, it is unclear how such consistency at organ scale can emerge from cellular behavior. We examine the growth of cell lineages, or groups of cells that are the progeny of a single mother cell. At early stages of the lineage, we find that initially smaller lineages grow faster than the larger ones reducing variability in lineage size, a phenomenon we refer to as size uniformization. In contrast at later stages of the lineage, size variability is enhanced when initially larger cell lineages grow faster than the smaller ones. Our results imply that the cell lineage changes its growth pattern at a tipping point. Finally, we found that the growth heterogeneity of individual cells within a lineage is correlated with fast growth of the lineage. Consequently, fast growing lineages show greater cell growth heterogeneity, leading to uniformization in lineage size. Thus, cellular variability in growth contributes toward decreasing variability of cell lineages throughout the sepal.

## Introduction

As in most living multicellular organisms, plant organs are reproducible; organs have their own characteristic sizes/shapes, making them landmarks for species identification in botany. In a naive sense, we might expect reproducible organs to arise from uniform cells with constant sizes and shapes like tiles in a floor. However, live imaging demonstrates that in many cases cell sizes, growth rates and directions exhibit considerable variability (Roeder et al., 2010, 2012; Hong et al., 2016; Elsner et al., 2012, Kierzkowski et al., 2012, Tauriello et al., 2015; Uyttewaal et al., 2012). For instance, the timing and geometry of cell division is variable in organs with stereotypical shapes (Roeder et al., 2010, Besson and Dumais, 2011). Those results raise the question of the contributions of such cellular noise and variability to organ size/shape consistency (Meyer and Roeder, 2014; Hong et al., 2016).

The complexity of the relationship between cells and organs can be illustrated with a few famous and mysterious biological examples. The first is the “regulative egg”; in specific early stages, when half of the early *Xenopus* embryo is removed, the remaining half produces a complete tadpole with half organ size (Spemann et al., 1924, Cooke et al., 1975). This suggests that the cell lineage is not the main driver for final size and shape, and that cell fate can be determined by the relative location within the embryo. In that scenario, cells would not be fully autonomous but instead subordinate to the whole shape and function of the embryo. A second example is “compensation”; when a mutation inhibits cell division and consequently reduces the number of cells in the organ, individual cells compensate that loss by increasing their size to produce an organ with nearly the correct size and shape (Tsukaya et al., 2003). This phenomenon of compensation suggests that organs have a global size/shape-sensing mechanism, which makes cell growth subordinate to the whole organ size/shape. Yet, as mentioned above, cells retain an ability to display variable growth rates, which suggests that cells are also autonomous to a large extent (Asl et al., 2011, Elsner et al., 2012). Therefore, we are left with a picture in which development results from a balance between the organismal theory (Kaplan et al., 1991; cell behavior is the consequence of the organ behavior) and the cell theory (organ behavior is the consequence of cell behavior). To shed light on the mechanisms balancing individual and collective behaviors in cell growth, we chose to focus on an intermediate scale, the group of cells, using a kinematic approach.

Here we focus on cell lineages (i.e. group of cells that descend from the same mother cell progenitor) of *Arabidopsis* sepals as an attempt to identify a unifying mechanism, which would also be compatible with both the cell theory and the organismal theory. Interestingly, Tauriello et al. used a kinematic approach to extract the growth of the cells with a same lineage (denoted as cell lineage group below) in order to determine general properties of the growth curves (Tauriello et al., 2015). Surprisingly, they found that the sizes of different cell lineage groups follow the same sigmoidal function of time, albeit with a stochastic timing of maximal growth rate, implying that the cell lineage groups do not grow “freely” but instead are constrained. Since these growth curves start from different initial cell sizes, the exact contribution of initial size distribution in such growth patterns becomes a central question. We investigated in this study the detailed kinematics and relations between the growth behaviors and the starting size of cell lineage groups of *Arabidopsis* sepals.

## Results

### 1. Switch from size uniformization to size variability enhancement in cell lineage groups

First, we investigated the relation between the initial sizes of the cells and their growth to determine whether uniformization occurs (e.g. smaller cells grow faster than larger cells) in developing *Arabidopsis* sepals. Live imaging data from two laboratories (5 wild-type sepals) previously reported in Hervieux et al., 2016 were considered. In this study, cells in the sepals outlined with plasma membrane markers were imaged every 12 or 24 hours. Since the size of individual cells changes after cell divisions, we considered the growth of cell lineage groups, i.e., the outermost cell contour inherited from the initial cell ignoring newly built anticlinal cell walls. The effect of growth of individual cells will be discussed in the Section 6. To extract the outline and follow the growth of cell lineage groups, we used analysis and visualization software “MorphoGraphX” (Barbier de Reuille et al., 2015; see Material and Method) for cell segmentation, lineage tracking and cell area calculations. We defined the cell area enclosed by the outline of a cell lineage group at time *t* as *A*_*t*_ and calculate the relative areal growth of the cell lineage group as (*A*_*t*+Δ*t*_ − *A*_*t*_)/*A*_*t*_ × 100 (%) (Fig. 1). The growth patterns in all analyzed sepals are qualitatively similar (Fig. S1.1-S1.4).

**Fig. 1.**
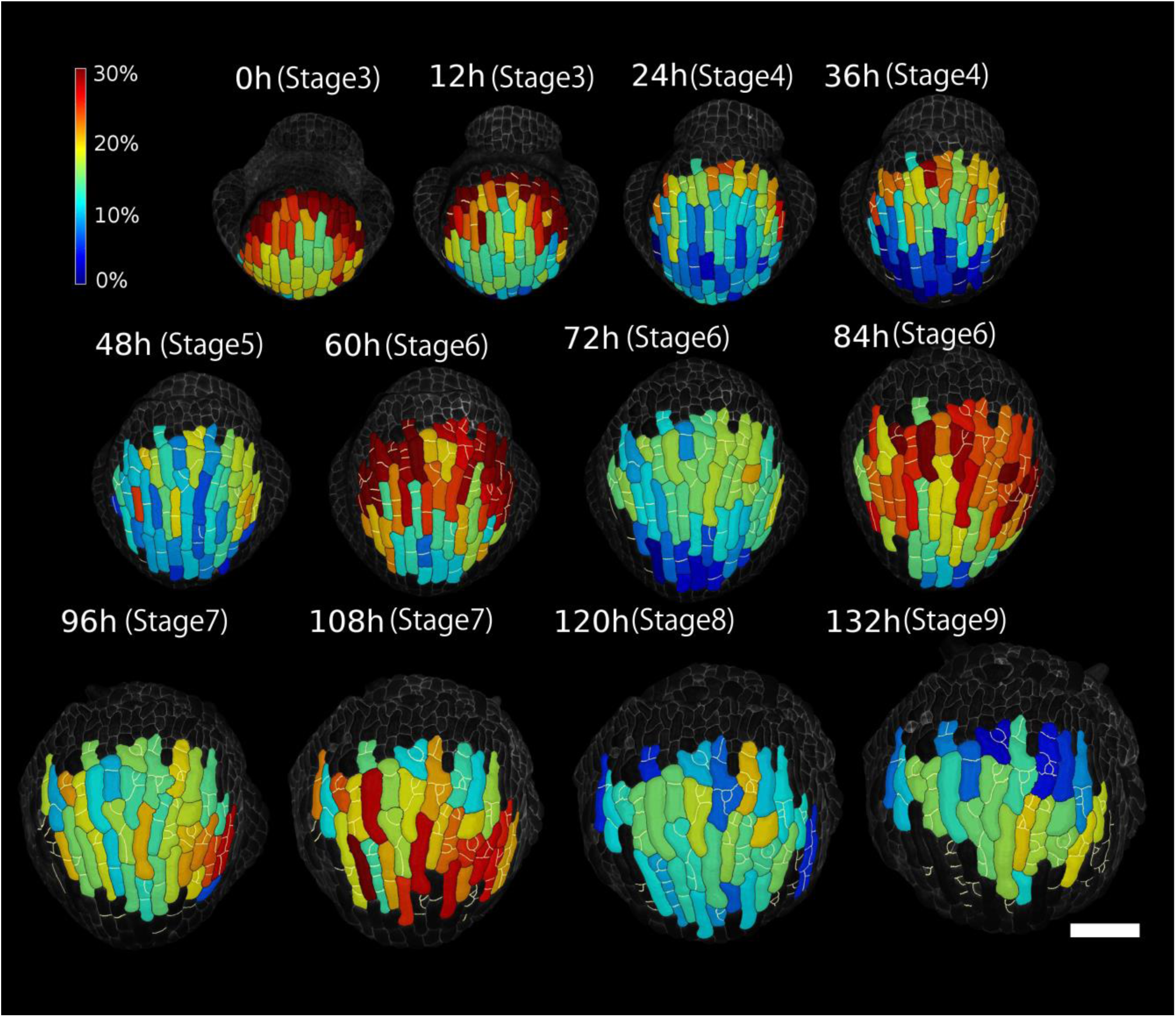
Spatio-temporal growth pattern of cell lineage groups. Heat map of areal growth of cell lineage groups (*A*_*t*+Δ*t*_ − *A*_*t*_)/*A*_*t*_ × 100 (%) over consecutive 12*h* intervals for flower wt-a1. The cell lineage groups and the new cell walls built after 0h are outlined in black and white, respectively. Note that around flower stage 5 to 6, there is a higher growth rate at nighttime and a relatively lower growth rate at daytime. Scale bar is 50*μm*.

As reported in our previous study (Hervieux et al., 2016), the sepal first undergoes fast growth at the tip (stage 3-4 in Fig. 1). The maximal growth then gradually moves down to the middle (stage 5-7) and bottom of the sepal (stage 8-9) in Fig. 1 (see also Fig. S1.1-S1.4). The general growth trend in area of the cell lineages in each sepal can be captured by the average of *A*_*t*_ (Fig. 2A). Although sepals from different laboratories (wt-a and wt-b shown in Fig. 2A) are slightly different because of different plant culture conditions, sepals within a given laboratory display comparable growth curves. In Fig. 2 and Table 1, we provided the flower stages of the sepals determined based on Smyth et al., 1990 (see also Materials and Methods).

**Table 1.** Flower stages of the sepals.

|  | wt-a1 | wt-a2 | wt-b1 | wt-b2 | wt-b3 |
| --- | --- | --- | --- | --- | --- |
| 0h | S3 | S3 |  |  |  |
| 24h | S4 | S4 |  | S3 |  |
| 48h | S5 | S5 | S5 | S4 | S6 |
| 72h | S6 | S6 | S6 | S6 | S7 |
| 96h | S7 | S7 | S7 | S7 | S7 |
| 120h | S8 | S8 | S8 | S8 | S8 |
| 144h | S9 | S9 | S9 | S9 | S9 |
| 168h |  |  | S9 | S9 | S9 |
| 192h |  |  |  | S10 |  |

**Fig. 2.**
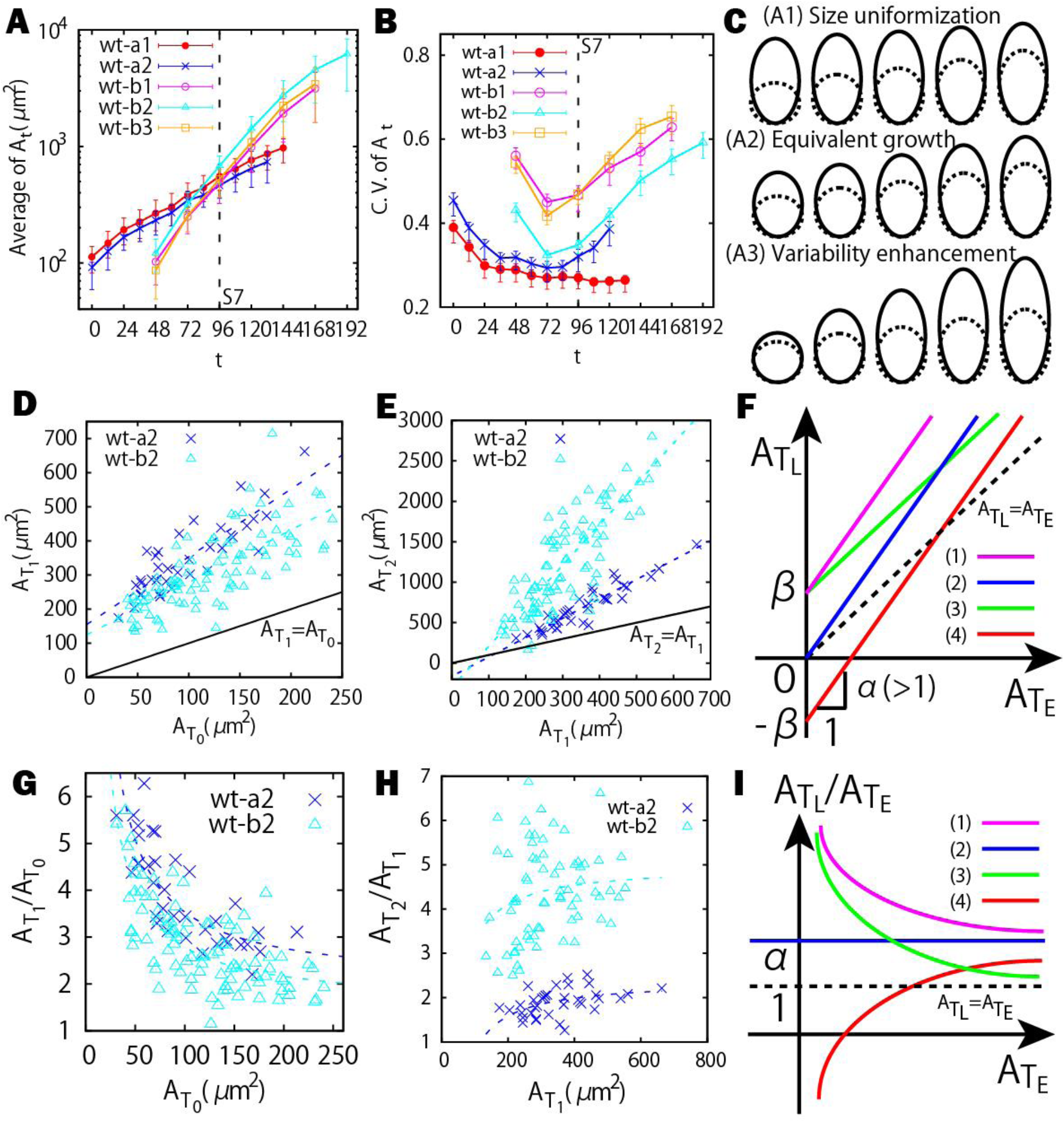
Size uniformization and variability enhancement in cell lineage groups. (A) The average area of the cell lineage groups in time. (B) Coefficient of variation (CV) to quantify area variability. Error bars represent the 50%-confidence interval (see Materials and Methods). CV decreases first and increases later. Note that the sepal wt-a1 does not show an obvious increase which could be due to the fact that initial cells in wt-a1 were imaged mainly from the tip to middle part of the sepal. (C) Schematic illustration of the three growth trends with earlier group size (dotted lines) and later group size (solid lines): (A1) size unifomization, initially smaller lineage groups grow faster, (A2) equivalent growth, lineage groups grow independent of their initial sizes, (A3) variability enhancement, initially larger groups grow faster. (D-E) 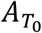 versus 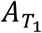 plot for the cell lineage groups (D), and 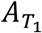 versus 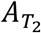 plot (E). The dashed lines are linear fittings of the data points in the plot of 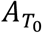 versus 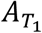. (F) Examples of different growth trends showing linear correlation between the area at a later time 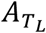 and those at an earlier time 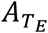. The green (1) and pink (3) curves illustrate uniformization, the blue (2) curve illustrates equivalent growth, and the red (4) curve illustrates variability enhancement. (G-H) 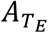 versus 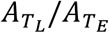 plot for the cell lineage groups at *T*_0_ versus those at *T*_1_ (G), and at *T*_1_ versus those at *T*_2_ (H). The dashed lines in (G-H) are fittings of the data points corresponding to those of (D-E), respectively. (I) Examples of different growth trends between 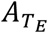 and 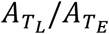. The line colors are the same as in (F). Size uniformization (variability enhancement) corresponds to the negatively (positively) correlated case.

Next, we analyzed the uniformization of the cell lineage groups, i.e. relating initial cell sizes to the growth of the associated cell lineage groups. To do so, we calculated the coefficient of variation (CV defined as standard deviation divided by mean) of *A*_*t*_ as a function of *t* (Fig. 2B). There are three possible trends in the area variability of the cell lineage groups quantified by the CV (Fig. 2C): (A1) Size uniformization during which initially smaller cell lineage groups grow faster to catch up with the larger groups (upper panel), leading to a decrease of CV; (A2) Equivalent growth in which the lineage groups grow homogeneously and independently of their initial sizes (middle panel), leading to an unchanged of CV; (A3) Variability enhancement during which initially larger cell lineage groups grow faster to outrun the smaller groups, leading to an increase of CV (lower panel). We observed a consistent behavior in the form of a decrease followed by an increase of CV, indicating that the growth of cell lineage groups undergoes a stereotypical transition in its cellular variability during the organ development.

To further analyze the switch from size uniformization to variability enhancement, we next determined how the area of each cell lineage measured at an early time relates to its area at a later time. We performed this comparison both during the size uniformization stage and the variability enhancement stage. To do so, we first defined some relevant times *T*_0_, *T*_1_ and *T*_2_ in our analysis. *T*_0_ denotes the starting time of our live imaging data which is in between stage 3 to stage 6 (see Table 1). For simplicity, *T*_0_ is set to be 0h for the sepals wt-a1 and -a2 that were imaged from the earliest stages among the samples (Table 1 and 2). *T*_1_ denotes the time when the CV reaches its local minimum, i.e., the tipping point. *T*_2_ denotes a later time point after *T*_1_ that is typically around flower stage 8 (see Table 1 and 2). During the size uniformization (A1) period [*T*_0_, *T*_1_], we observe a linear correlation between the cell lineage area at 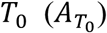 and the cell lineage area at 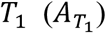 (Fig. 2D). Likewise, we observe a linear correlation between cell lineage area at 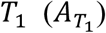 and the cell lineage area at 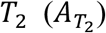 during the variability enhancement (A3) period [*T*_1_, *T*_2_] (Fig. 2E).

**Table 2.** Fitting results for the linear relation 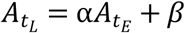 for the period [*T*_*E*_, *T*_*L*_] = [*T*_0_, *T*_1_] and the period [*T*_*E*_, *T*_*L*_] = [*T*_1_, *T*_2_]. R^2^ is the coefficient of determination.

| | Period $[T_0, T_1]$: $A_{T_1} = \alpha_1 A_{T_0} + \beta_1$ | | | | |
| --- | --- | --- | --- | --- | --- |
| | $T_0(h)$ | $T_1(h)$ | $\alpha_1$ | $\beta_1$ | $R^2$ |
| Wt-a1 | 0 | 108 | $2.37 \pm 0.51$ | $338 \pm 65.2$ | 0.463 |
| Wt-a2 | 0 | 72 | $1.99 \pm 0.20$ | $155 \pm 22.1$ | 0.741 |
| Wt-b1 | 48 | 72 | $1.81 \pm 0.07$ | $63.4 \pm 7.61$ | 0.838 |
| Wt-b2 | 48 | 72 | $1.54 \pm 0.14$ | $125 \pm 18.7$ | 0.564 |
| Wt-b3 | 48 | 72 | $1.37 \pm 0.16$ | $110 \pm 15.2$ | 0.330 |
| | Period $[T_1, T_2]$: $A_{T_2} = \alpha_2 A_{T_1} + \beta_2$ | | | | |
| | $T_1(h)$ | $T_2(h)$ | $\alpha_2$ | $\beta_2$ | $R^2$ |
| Wt-a1 | 108 | 132 | $1.34 \pm 0.06$ | $11.8 \pm 41.4$ | 0.945 |
| Wt-a2 | 72 | 120 | $2.39 \pm 0.16$ | $-163 \pm 57.9$ | 0.872 |
| Wt-b1 | 72 | 120 | $4.03 \pm 0.23$ | $-48.0 \pm 59.8$ | 0.712 |
| Wt-b2 | 72 | 120 | $5.02 \pm 0.38$ | $-295 \pm 122$ | 0.673 |
| Wt-b3 | 72 | 120 | $5.03 \pm 0.26$ | $-238 \pm 67.25$ | 0.748 |

Since we observed linear correlations in 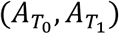 and 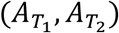 (Fig. 2D and 2E), we suspect that the key parameter for the transition between size uniformization and variability enhancement could be embedded in a linear equation relating lineage area at the earlier time 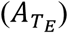 to lineage area at the later time 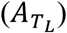, i.e., 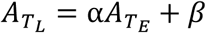 (Fig. 2F). Since almost all cell lineage groups increase their areas from *T*_*E*_ to *T*_*L*_, the parameter α is greater than or equal to 1. Under this setup, the cumulative growth ratio (i.e. final lineage size divided by initial lineage size), defined as 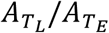, can be easily derived as 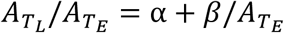. In Fig. 2G and 2H, we illustrate different growth behaviors in the plots of 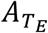 versus 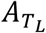 and 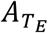 versus 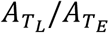. As shown in Fig. 2I, the size uniformization case (A1) (initially smaller cells grow faster i.e., 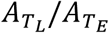 negatively correlates with 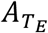) occurs when β > 0 (e.g., pink curve labeled as (1) and the green curve labeled as (3)). The equivalent growth case (A2) (cell growth does not depend on cell size i.e., 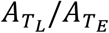 is independent of 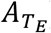) corresponds to β = 0 (blue curve labeled as (2)). The variability enhancement case (A3) (initially larger cells grow faster i.e. 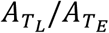 positively correlates with 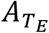) occurs when β < 0 (e.g. red curve labeled as (4)).

Size uniformization (A1) was quantitatively confirmed before the tipping point *T*_1_ from the growth behavior during the period [ *T*_0_, *T*_1_] (Fig. 2G), and variability enhancement (A3) was confirmed after *T*_1_ from the growth behavior in the period [*T*_1_, *T*_2_] (Fig. 2H). The fitted parameters before the tipping point (α_1_, β_1_) satisfy α_1_ > 0 and β_1_ > 0 for uniformization (Table 2). On the other hand, after the tipping point (α_2_, β_2_) satisfy α_2_ > 0 and β_2_ < 0 for the sepals wt-a2, wt-b1, wt-b2, wt-b3 consistent with variability enhancement. However, β_2_ > 0 for the sepal wt-a1 because wt-a1 shows more or less the case (A1) or (A2) (Fig. S2D) consistent with the CV analysis showing that wt-a1 remains in uniformization and does not reach the tipping point (Fig. 2B). To summarize, we concluded that a size uniformization mechanism is reducing the cell lineage area variability initially before the lineages switch to variability enhancement (Fig. 2B), demonstrating the presence of a transition of growth behavior during sepal development. In addition, we find that the parameter β corresponding to the y-intercept in the plot of 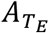 versus 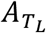 (see Fig. 2F) provides us with an effective parameter for identifying such a growth transition.

### 2. Size uniformization occurs everywhere in sepal

One of the implications from the linear positive correlation between earlier lineage group sizes and the later ones in both the size uniformization and variability enhancement (Fig. 2D and 2E) is that the rank of the cell lineage group sizes is largely preserved during sepal development. This means that although smaller lineage groups grow faster in the size uniformization stages, their growth rates are not fast enough to catch up with, or even outrun, the initially larger groups. We suspect that some kinds of spatial information of the lineages play a role in the ordering of their size. To verify this, we track the areas of the cell lineage groups *A*_*t*_ in both space and time during the size uniformization stages (Fig. 3A and 3B) by coloring the growth curves in *A*_*t*_ according to their initial size at time *T*_0_ (Fig. 3B). The color gradient at earlier times is largely preserved at later times (Fig. 3B and Fig. S3), confirming again that the size order of the cell lineage groups is largely preserved. At the initial time *T*_0_, smaller cells are distributed toward the top part of the sepal and the larger ones toward the bottom. This is because the cells in the top part are farther along in their development and have started dividing at the early flower stages. As expected, the spatial distribution of size is not strongly affected by growth and the cells at the tip remain smaller than those at the bottom of the sepal at a later time *T*_1_ (Fig. 3C and Fig. S3).

**Fig. 3.**
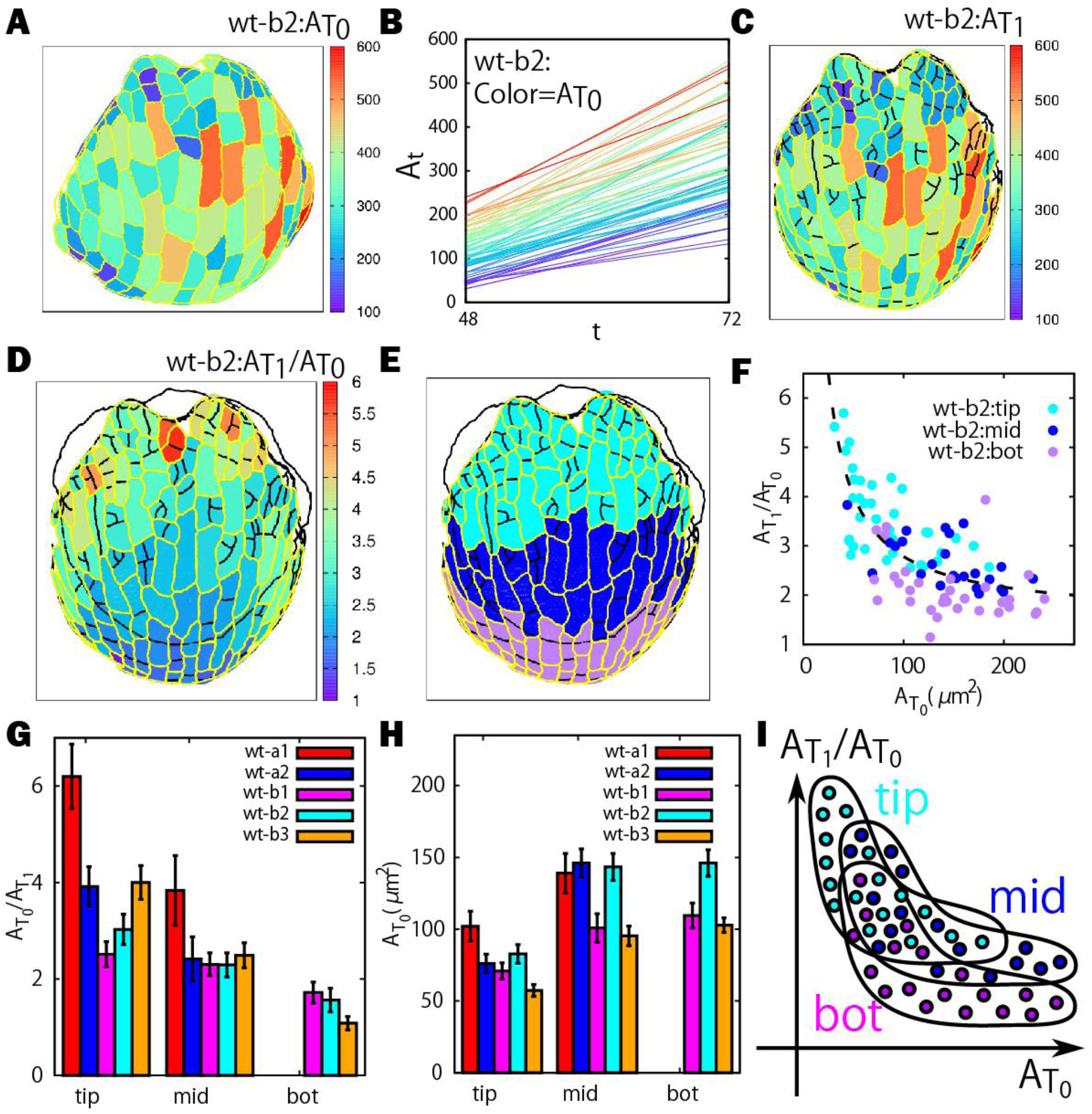
Size uniformization occurs everywhere in sepal. (A) Heat map of the initial cell area 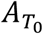. (B) Growth curves of cell lineage groups colored according to their initial size 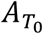. (C) Heat map of the size of the cell lineage group 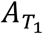 at a later time *T*_1_. (D) Heat map of the cumulative growth ratio 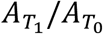 showing a continuous spatial trend from tip to bottom. (E) Classification of tip (cyan), middle (blue) and bottom (purple) regions. (F) Plot of cumulative growth ratios 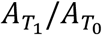 versus initial area 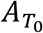 for cell lineage groups in each region. (G-H) Histograms of cumulative growth ratios (G) and initial area (H), for the three regions. (I) Schematic illustration of positional dependence of size and growth.

Next, we looked at the spatial distribution of the cumulative growth ratio 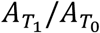 on the sepal (Fig. 3D). There is a continuous change from the high values in 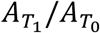 at the tip part to the lower values at the bottom part for all the sepals we analyzed. To investigate whether the size uniformization occurs only in specific regions (e.g. tip) of the sepal, we manually divided the sepal into the tip, middle and bottom regions as indicated by cyan, blue and purple colors, respectively, (Fig. 3E and Fig. S4). 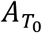 negatively correlated with 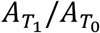 for cell lineage groups from each region showing that size uniformization occurs throughout the sepal (Fig. 3F and Fig. S4). The plots from different regions were slightly shifted relative to one another as expected given the overall trends of growth in the sepal, namely, tip cells have smaller 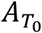 but larger 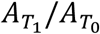, whereas bottom cells have larger 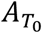 but smaller 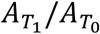 (see also the histograms in Fig. 3G and 3H). Although the distributions of 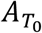 and 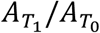 are slightly different as schematically illustrated in Fig. 3I, our results indicate that size uniformization occurs as a global mechanism in the developing sepal and the spatial information does not play a significant role in regulating the size of the cell lineage groups during the size uniformization stages. During the variability enhancement stages, we could not find out a characteristic trend in Fig. S5.

### 3. Temporal variation of correlation between initial size and growth at each time step

As the size uniformization does not depend on the location of cell lineage groups, we then asked whether there is gradual or sudden temporal change in the correlation between the initial lineage group size 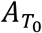 and the growth ratio (*A*_*t*+Δ*t*_/*A*_*t*_)/*Δt*. In other words, we examined the correlation in short time intervals where *Δt* is 12 or 24 hours (compared with the long time intervals analyzed in the previous sections). The initial sizes of cell lineage groups are negatively correlated with the growth ratios during the first 0-72h (or stage 4-7) whereas at later stages the correlations become positive (Fig. 4A). The change of correlations from negative to positive was further confirmed by calculating, at each time step, the Spearman correlation coefficient (SCC) between the initial sizes 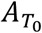 and the growth ratios (*A*_*t*+Δ*t*_/*A*_*t*_)/*Δt* from all cell lineage groups. The SCCs indicate that the growth ratio at each time step is negatively correlated at the earlier stages but becomes positively correlated at the later stages (Fig. 4B). The transition took place at around stage 6-7 with an exception that the SCC for the sepal wt-a1 has a time delay of ∼36h. These results also show that there is a temporal variation changing from negative correlation during the size compensation to positive correlation during the variability promotion stages as shown in Fig. 4B, meaning that the transition takes place gradually.

**Fig. 4.**
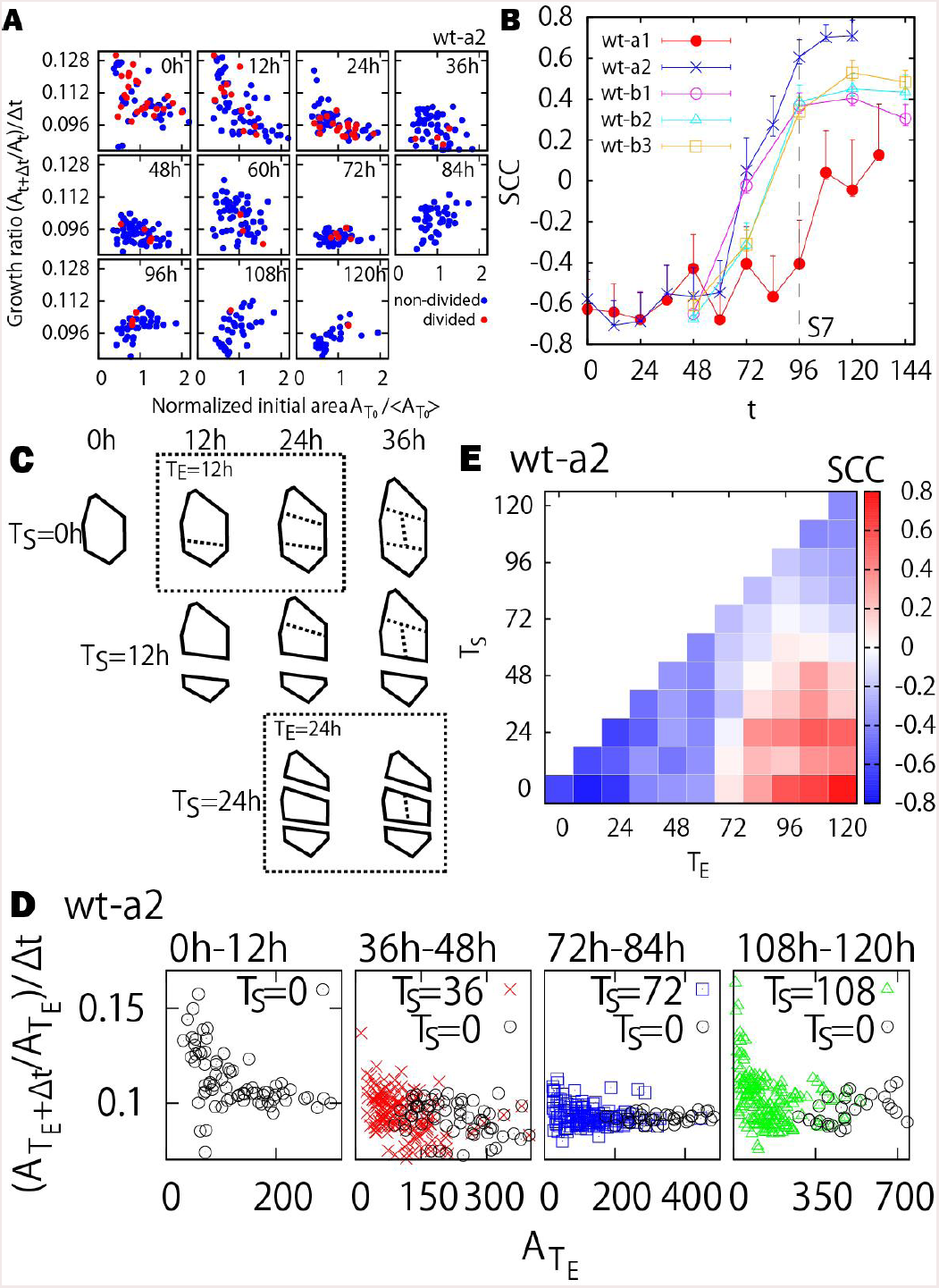
Size uniformization depends on starting time of lineage. (A) Plot of growth ratio at each time step (*A*_*t*+Δ*t*_/*A*_*t*_)/*Δt* versus normalized initial area 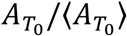 for the sepal wt-a2. Here the normalized initial area is considered for a better visual comparison of areas at different time points. The dividing and non-dividing lineage groups are colored in red and blue, respectively. (B) Spearman correlation coefficient (SCC) for the plots in (A). The result shows that the SCCs change from negative to positive at around stage 6-7 (72h-96h). (C) Schematic illustration of the definition of the cell lineage group from starting time T_S_. The dotted boxes show some examples of the definition (*T*_*E*_ = 12*h* for *T*_*S*_ = 0 or *T*_*E*_ = 24*h* for *T*_*S*_ = 24). (D) Diagram of size and growth rate for different cell lineage groups. (E) Spearman correlation coefficient for different combination of *T*_*E*_ and *T*_*S*_.

### 4. Shift from size uniformization to variability enhancement occurs around 24∼60 hours after a lineage initiates from a single cell

A remaining question is whether the switch from size uniformization to variability enhancement relates to the developmental timing of the entire sepal, or relates to the timing at which an individual cell lineage initiates from a single cell as our analysis in the previous sections started from single cells at the first time point of the live imaging series. To answer this question, we considered lineages starting from single cells at additional time points during the live imaging sequence of a given sepal, denoted as the starting time *T*_*S*_, at which the set of all individual cells is extracted with their cell walls and areas identified (Fig. 4C leftmost outlines). Each individual cell defined at *T*_*S*_ therefore serves as the mother cell of a particular cell lineage group for *t* > *T*_*S*_, i.e., a different choice of *T*_*S*_ (= 0h, 12h, 24h,…) results in different choice of cell lineage groups (Fig. 4C). The different choice of *T*_*S*_ is equivalent to different starting times (denoted by *t* = 0h in Table 1) of the live imaging, which can be arbitrary. We then asked if the signature of size uniformization, i.e., the negative correlations observed in Fig. 4A and 4B, depends on the choice of *T*_*S*_.

As previously (in Fig. 4A), we examined how the size of the cell lineage group 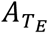 at *t* = *T*_*E*_ (with *T*_*E*_ > *T*_*S*_) correlates with the growth ratio 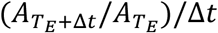 (with Δ*t* = 12h) at each time step (see e.g. the dash boxes in Fig. 4C for different choices of *T*_*E*_). Fig. 4D provides several examples of the plot 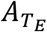 versus 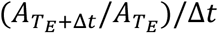 with different choice of *T*_*S*_ and *T*_*E*_. We see that whenever a new lineage is initiated from a single cell, it undergoes size uniformization, i.e. there is a negative correlation between cell size and growth rate indicating that smaller cell lineages grow faster than larger cell lineages (Fig. 4D). Simultaneously, a lineage initiated at an earlier time *T*_*S*_, may have already transitioned to variability enhancement. For example, the cell lineage initiated at 108 hours is undergoing size uniformization, while the cell lineage initiated at 0 hours is undergoing variability enhancement (Rightmost panel in Fig. 4D). Since the single cells initiating lineages at 108 hours are included within the 0-hour cell lineages, this constitutes an intricate multiscale system.

We verified that this transition relates to the starting time of the lineage with Spearman correlation coefficients for different *T*_*S*_ and *T*_*E*_ in which blue and red regions correspond to the size uniformization (A1) and variability enhancement (A3) cases, respectively (Fig. 4E and S6). Interestingly, the growth transition between (A1) and (A3) presents as *T*_*S*_ varies. Furthermore, the transition time from the blue to red regions at which the corresponding Spearman correlation coefficients (SCCs) in Fig. 4E show that the growth transition from uniformization to variability enhancement gradually shift to later time depending on the starting time T_S_. On average, the transition occurs ∼60 hours for wild type a (∼24 hours for wild type b in Fig. S6) after the initiation of the lineage from a single cell at *T*_*S*_. This means that the growth transition observed in Fig. 4A and 4B depends on the starting time of lineage in a way that the cell lineage groups have to grow up to a certain size (or scale) for variability enhancement to happen. We also emphasized that the transition does not depend on the overall developmental timing of the sepal.

### 5. Growth pattern of cell lineage group is independent of cell division and stomata differentiation

One of the other possibilities is that size uniformization could depend on the topography of cell lineage groups, e.g., on cell divisions and the presence of newly built walls. Furthermore, the tipping point *T*_1_ may also relate to the time when epidermal cells acquire a different fate, notably with the differentiation of stomata within cell lineage groups. To address this issue, we analyzed the correlation between the initial size 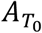 and the growth ratio at each time step (*A*_*t*+Δ*t*_/*A*_*t*_)/Δ*t* while keeping track of cell divisions in the cell lineage groups as shown by the colored dots in Fig. 4A. We defined the cell lineage groups as dividing at time *t* if cell division occurs within the lineage group during the time step (*t*, *t* + Δ*t*). The dividing lineage groups can also include cells that differentiate and divide into stomata. No significant difference between dividing and non-dividing cell lineage groups from the sepals can be observed (Fig. 4A and Fig. S7). Therefore, we concluded that the growth pattern of cell lineage groups is independent of cell divisions and differentiations.

### 6. Individual cell growth heterogeneity is positively correlated with growth at each time step in cell lineage groups

Since the cause of the tipping point cannot be simply related to a change in cell identity, we next investigated whether growth heterogeneity within the cell lineage group may be associated with the shift from uniformization to variability enhancement. To address this question, we quantified the degree of cell growth heterogeneity *G*_*h*_ at time *t*, that is defined as the CV of the cell growth ratio (*A*_*t*+Δ*t*_′/*A*_*t*_′)/*Δt* of all individual cells within a specific cell lineage group (see Materials and Methods). To determine the role of cell growth heterogeneity in size uniformization, we first asked if *G*_*h*_ correlates with the initial size 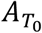 (Fig. 5A and 5B). We found that the correlation between growth heterogeneity and initial size is mostly negative at the early stages and then becomes positive at the later stages. In other words, a higher level of cell growth heterogeneity exists among the daughter cells for initially smaller cells at the size uniformization stages, while at the variability enhancement stages initially larger cells show higher growth heterogeneity (Fig. 5E).

**Fig. 5.**
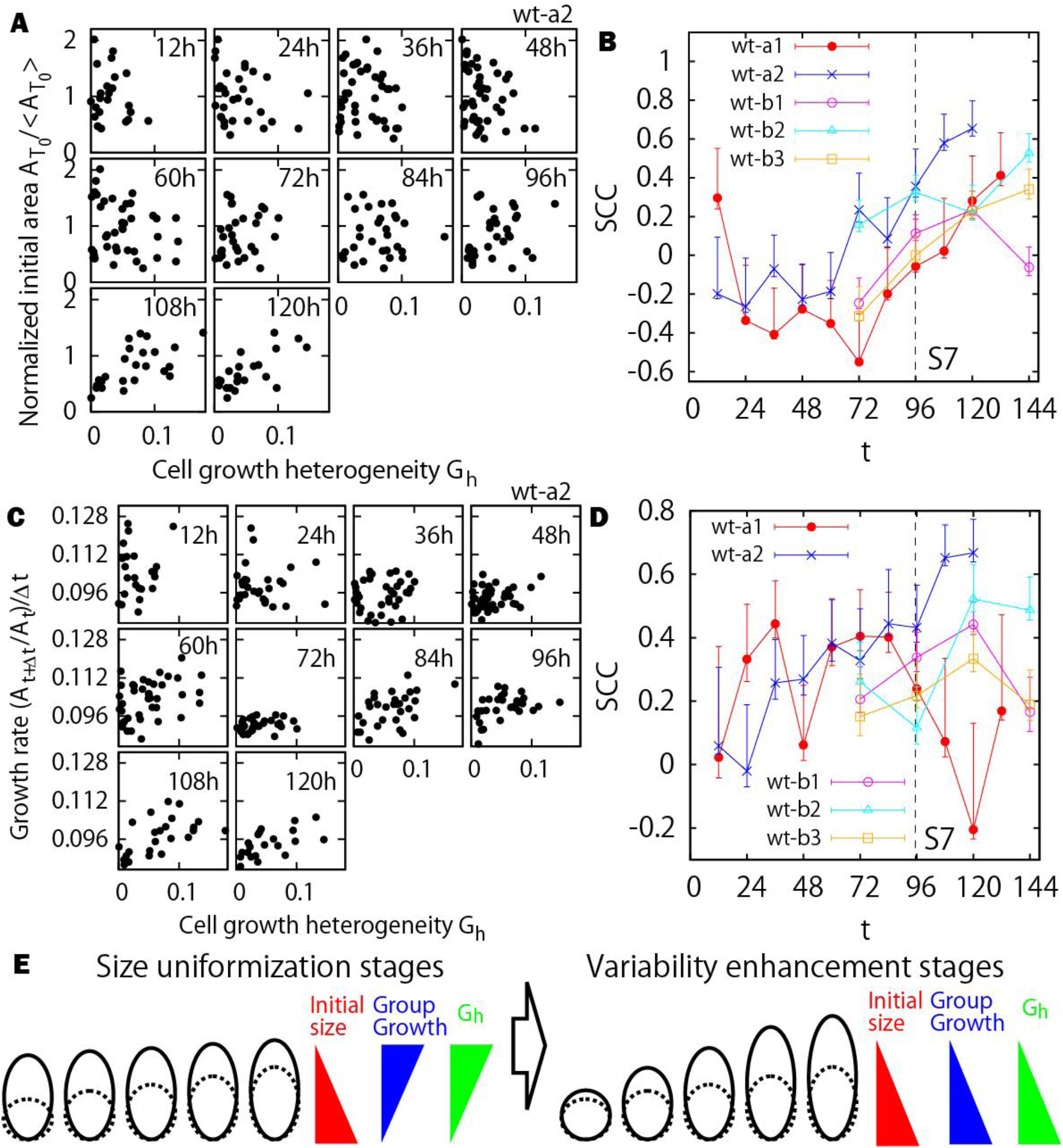
Increase of the cell growth heterogeneity and its correlation with initial size of the cell lineage group. (A) Cell growth heterogeneity versus normalized initial area. (B) Spearman correlation coefficient of (A). (C) Cell growth heterogeneity versus growth ratio of the cell lineage group at each time step. (D) Spearman correlation coefficient of (C). Note that the sepal wt-a1 has less correlation at the later stages that may relate to the time delay of the growth transition. (E) Schematic illustration of the results with earlier group size (dotted lines) and later group size (solid lines).

Based on the temporal change of correlation between initial size and *G*_*h*_ (Fig. 5B), and a similar one between the initial size and the growth ratio at each time step (Fig. 4B), a positive correlation between *G*_*h*_ and the growth ratios of the cell lineage groups at each time step is expected for both the size uniformization and variability enhancement stages. Such positive correlations were confirmed in Fig. 5C and 5D, namely, a faster growing cell lineage group during a time step of *Δt* (i.e., 12h in our case) has a larger growth heterogeneity (or variability) among individual cells belonging to group. These finding suggests that growth variability at the cellular level could enhance the growth of the cell lineage group. The results of the other sepals are qualitatively similar (Fig. S8). Alternatively, the fast growing groups could exhibit more heterogeneity in individual cell growth rates because they have to accommodate the slower growth of neighboring cell groups. This factor could play an important role in plant cells, since the cells are glued to their neighbors and cannot slide against each other like animal cells do. Finally, fast growing groups of cells are more likely to initiate stomata, which initially grow faster than their immediate neighbors and can therefore explain the increased growth heterogeneity within the group.

Altogether, these analyses demonstrate that a shift from size uniformization to variability enhancement occurs, which is correlated with a change in growth heterogeneity, rather than cell position or cell identity (Fig. 5E).

## Discussion

To summarize, our results reveal the existence of a growth phase transition between two growth modes in the developing sepal. The size uniformization mode occurs after the initiation of cell lineages where initially smaller cell lineage groups grow faster relative to the larger ones resulting in a decrease in the area variability of the lineage groups, and the variability enhancement mode occurs at the later stages of the lineages where initially larger cell lineage groups grow even faster resulting in an increase of area variability. While we exclude a role of cell position or cell identity in the shift from size uniformization to variability enhancement, we find that the size uniformization can be, in principle, observed in any cell lineages soon after it initiates from a single cell. One of the surprising observations was that heterogeneity in growth between the cells within the lineage correlates with the faster growth of the cell lineage.

The dependence of the size uniformization with starting time of lineage group can be interesting in terms of scale dependence because the size uniformization is observed to occur at smaller cellular scale until the lineage groups reach a larger size, i.e., supracellular scale, in 24∼60 hours at which a different growth mechanism of size variability enhancement starts to emerge. This means that the growth behavior at the smaller scale can be different from those at larger scale. To fully understand this multiscale phenomenon, one possible way is to see the growth with gradual change in spatial scale and extract the statistical law of growth ratio as a function of scale. This type of growth analysis should be a key investigation for understanding the growth at the supracellular level.

The conventional sense of the compensation in plant biology was the balance between cell division and cell growth that leads to consistent organ size in mutants or transgenic lines due to changes in cellular size to compensate for changes in the number of cells in the organ (Marshall et al, 2012, Tsukaya, 2003). Here we detect a same kind of compensation, i.e., size uniformization in the wild type at the cell lineage scale. Recently, size uniformization was observed within sister cells in a single cell cycle in shoot apical meristem (Willis et al., 2016). We note that in our case the size uniformization was observed beyond specific cell cycle but with a much longer time range that spans several flower stages.

Our findings of the long-ranged temporal correlations (Fig. 2) suggest that a cell lineage group still keeps a memory of its initial size, even after a long time. An attractive (but still speculative) hypothesis is that the cell lineage group contour (i.e., the initial walls, including bottom and top walls) provides long-term cues to growth because they are inherited. Thus, it seems that a cell lineage group would behave like an individual big cell and act as a small organism that remembers its initial size.

The observed correlation between group growth and growth heterogeneity implies that the shift rather relies on perception of growth-related parameters. This raises the question of the exact nature of that cue. Growth heterogeneity generates mechanical conflicts (Rebocho et al., 2016) and plant cells are able to respond to such mechanical signals (Hervieux et al., 2016, Uyttewaal et al., 2012). Therefore, our analysis may be consistent with a scenario in which a competence to respond to mechanical cues is regulated in time, making unifomization possible during sepal development.

## Materials and Methods

### Plant material and imaging by confocal microscope

The five wild-type sepals are sampled from different laboratory (wild type a1, a2: two sepals imaged every 12h at ENS Lyon in France, wild type b1, b2, b3: three sepals imaged every 24h at Max Planck Cologne in Germany). Here we reanalyzed data already used in (Hervieux et al., 2016), where the average sepal growth pattern was quantified.

In a1 and a2 series, we used the *pUBQ10::myrYFP* line kindly provided by Raymond Wightman. In this line, myrYFP corresponds to a YFP that is N-terminally modified with a short peptide that is myristoylated and probably acylated (Yang et al., 2016). Plants were grown under long day conditions. Staging was determined as indicated in Smyth *et al.,* 1990. One- to two-cm-long main inflorescence stems were cut from the plant. To access young buds, the first 10–15 flowers were dissected. The young buds were imaged with an SP8 laser-scanning confocal microscope (Leica) using long-distance 40× (NA 0.8) water-dipping objectives. During time-lapse imaging, plants were kept in one-half MS medium with PPM (Plant protective medium) (1ml/L) and imaged every 24 hr for up to 8 days.

In the b1, b2 and b3 series, early stages of floral development (from 3 to 6) were determined based on comparison of floral bud morphology with the stages proposed by Smyth et al. (1990). The timing of development from stage 3 to 6 corresponded to the one published (Smyth et al. 1990) for similar growth conditions. Therefore, later developmental stages of the same flowers were determined by comparing the timing from stage 6 (bud closure) with the timing of development reported by Smyth and coworkers.

### Extraction of cell surface and cell growth extraction using software “MorphoGraphX”

To investigate the cell growth, we used an open source application for the visualization and analysis of a fluorescence data “MorphoGraphX (MGX)” (Barbier de Reuille et al., 2015). As described in Ref. 19, A key strength of MGX is the ability to summarize 3D fluorescence data as a curved surface image. From a fluorescence data, the MGX allows us to detect the outermost surface indicating the 3D organ shape. After creating the surface mesh which is the aggregation of small triangles along with the organ surface, we can also specify the position of cell walls (cell outlines in the main text) using a membrane fluorescence which has a strong brightness at the cell walls. Then, we can calculate a specific cell surface area to sum up the total area of small triangles within a cell. We also have a cell lineage information which combine a dataset observed at different times. In this paper, we first detected a cell outline at the first time frame and keep tracked of the outline in which cells have a same lineage as the initial cell. The growth in cell lineage group can be calculated as the total cell area of a same lineage at time *t* over the total cell area of a same lineage at time *t*, i.e. *A*_*t*+ *Δt*_/*A*_*t*_.

### Bootstrap method to estimate first and third quartiles of C. V

We calculated the first and third quartiles of CV using the bootstrap method. The bootstrap method can be summarized as “random draw with replacement” which allows us to measure the accuracy of the statistics. For the sake of a general description, suppose we have an observed valuable X (the sample number is N) and its CV (standard deviation of X / mean of X). We first create a bunch of “bootstrapped data set X^B^”. That is, we randomly drew samples N times from data set X with replacement and set it as X^B-1^ for the first trial. We keep doing the same way and create the data sets (X^B-1^, X^B-2^,…, X^B-1000^). Using this data set, we can get a bunch of “bootstrapped CV” (CV^B-1^, CV^B-2^,…, CV^B-1000^). We then calculated the first and third quartiles of the distribution of the bootstrapped CV.

### Quantification of cell growth heterogeneity

The cell growth heterogeneity *G*_*h*_ at time *t* is calculated by the CV of the individual cell growth 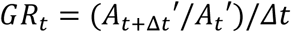 within a specific cell lineage group where 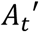 is the surface area of the individual cell at time *t*. Suppose we have M number of the individual cell growth rates, *GR*_*t*_(*C*_1_), at time *t* within a specific lineage group *C*_1_, we have the CV *CV*_1_(*C*_1_) for the specific cell lineage group data set 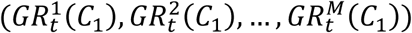. Likewise, if we applied this process to *K* number of cell lineage groups, we get the data set of the CV (*CV*_1_(*C*_1_), *CV*_2_(*C*_2_),…, *CV*_*K*_(*C*_*K*_)). Then the term “cell growth heterogeneity” in the main text is defined as the averaged value of the data set (*CV*_1_(*C*_1_), *CV*_2_(*C*_2_),…, *CV*_*K*_(*C*_*K*_)).

## Author contributions

S.T. and C-B.L. designed the work and drafted the manuscript with input from all the co-authors. N.H. and D.K. did the live imaging, A-L.R-K, A.S. and N.H. analyzed the 3D images to recreate the cells on the organ surface. All co-authors read and approved the final manuscript.

## Acknowledgements

We thank Mathilde Dumond, Lilan Hong, and Mingyuan Zhu for constructive discussions on this work.

## Competing interests

No competing interests declared.

## Funding

This work was supported by the Human Frontier Science Program grant RGP0008/2013 (A.B./O.H., A.H. K.R., C-B.L., R.S.S.) and a Special Postdoctoral Researcher Program in RIKEN (Japan).

---

**Fig. S1.1-S1.4.**
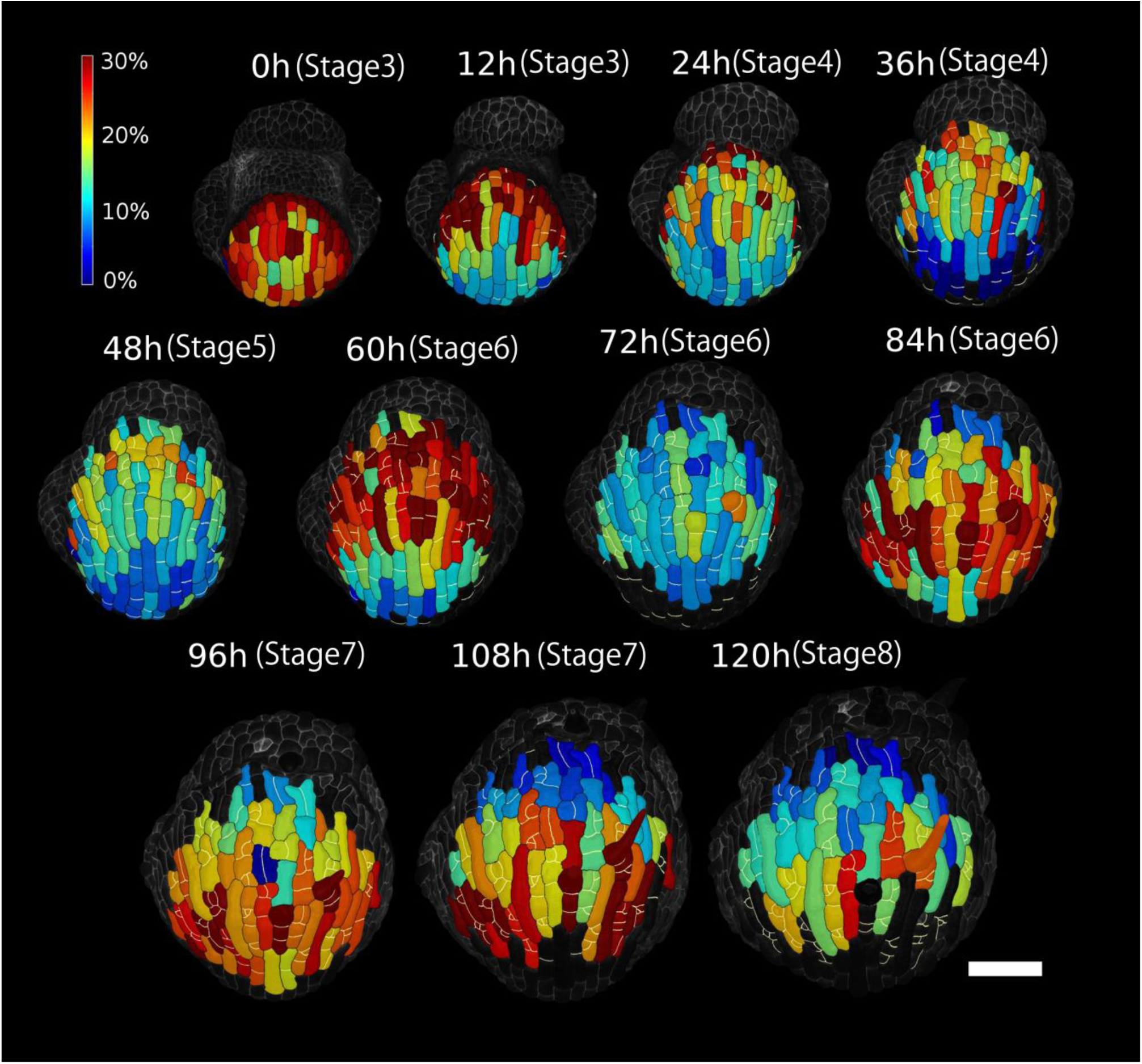
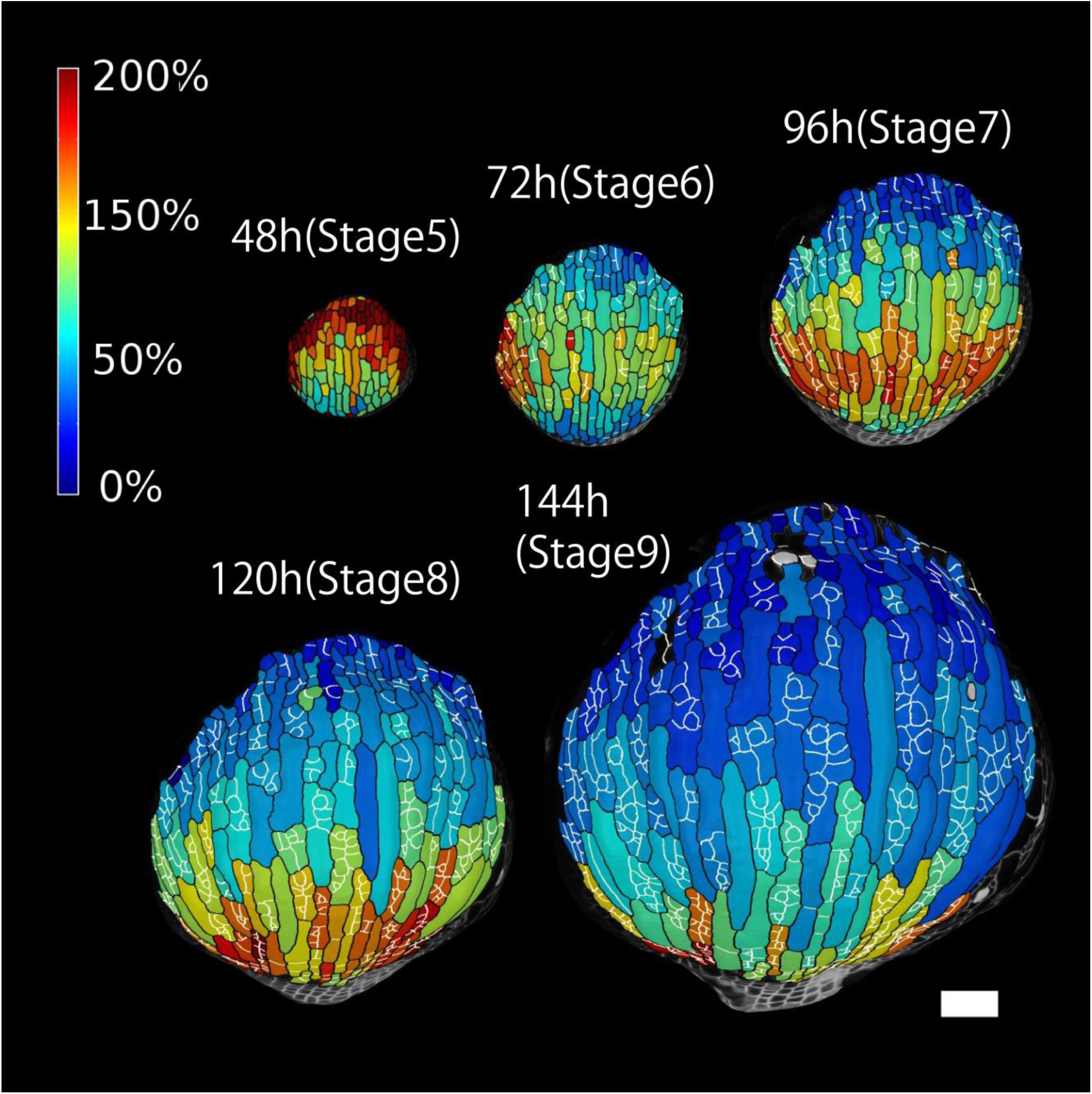
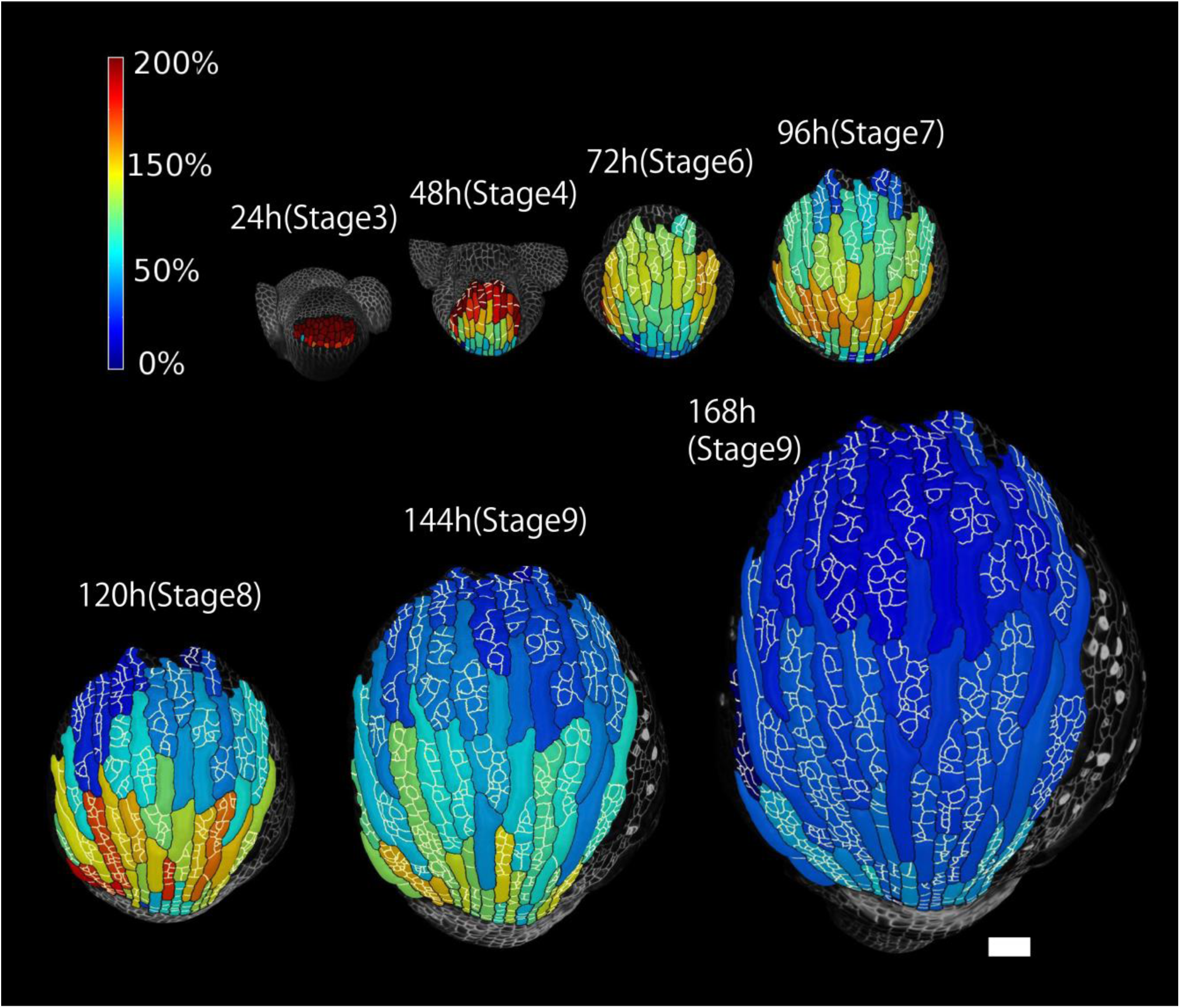
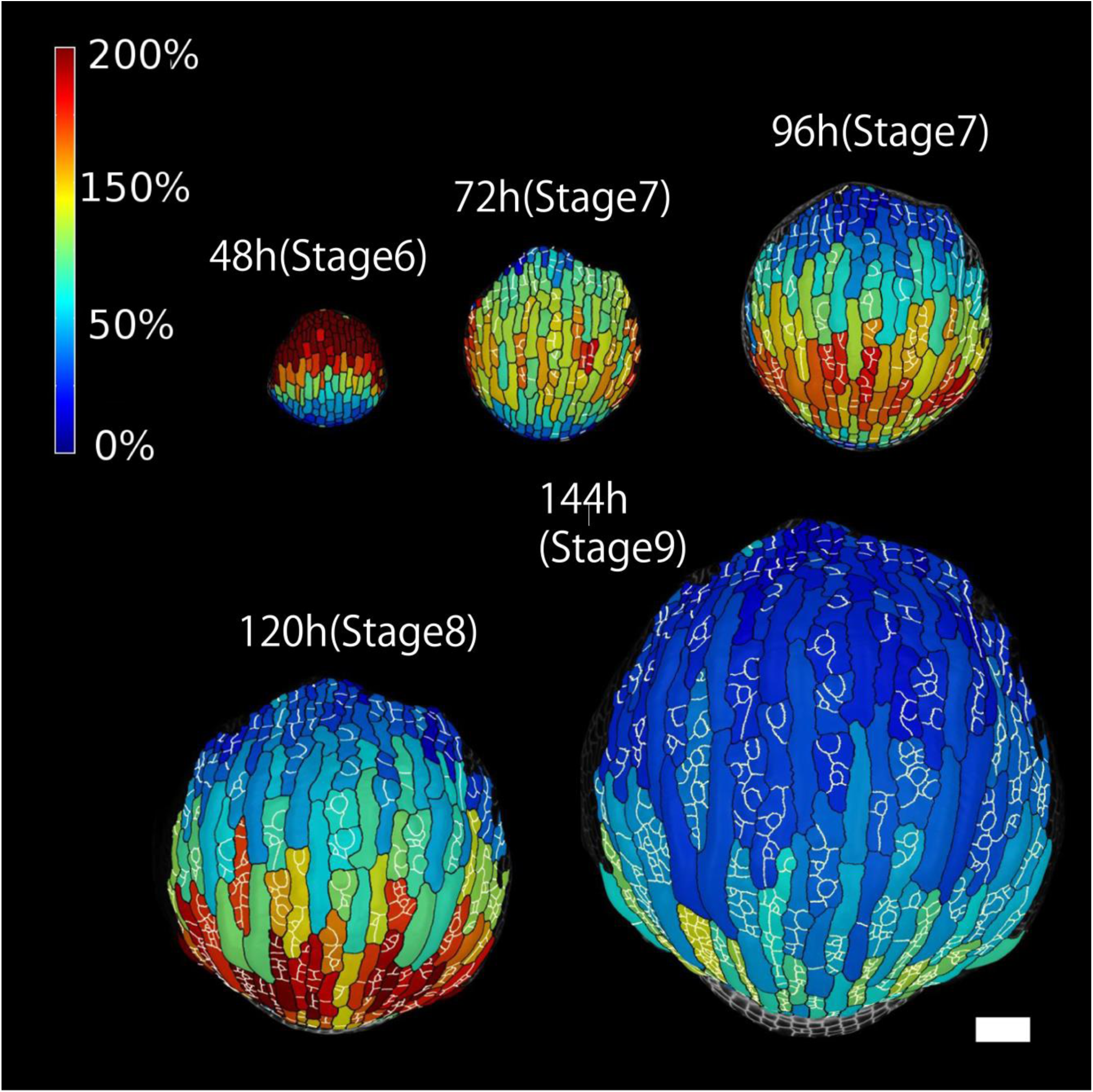
Heat map of areal growth of cell lineage groups (*A*_*t*+Δ*t*_ − *A*_*t*_)/*A*_*t*_ × 100 (%) over consecutive 12*h* intervals for flower wt-a2 (S1.1), and consecutive 24*h* intervals for flower wt-b1, wt-b2 and wt-b3 (S1.2-S1.4). Scale bars are 50*μm*. The cell lineage groups and the new cell walls built after 0h are outlined in black and white, respectively.

**Fig. S2. (A-B).**
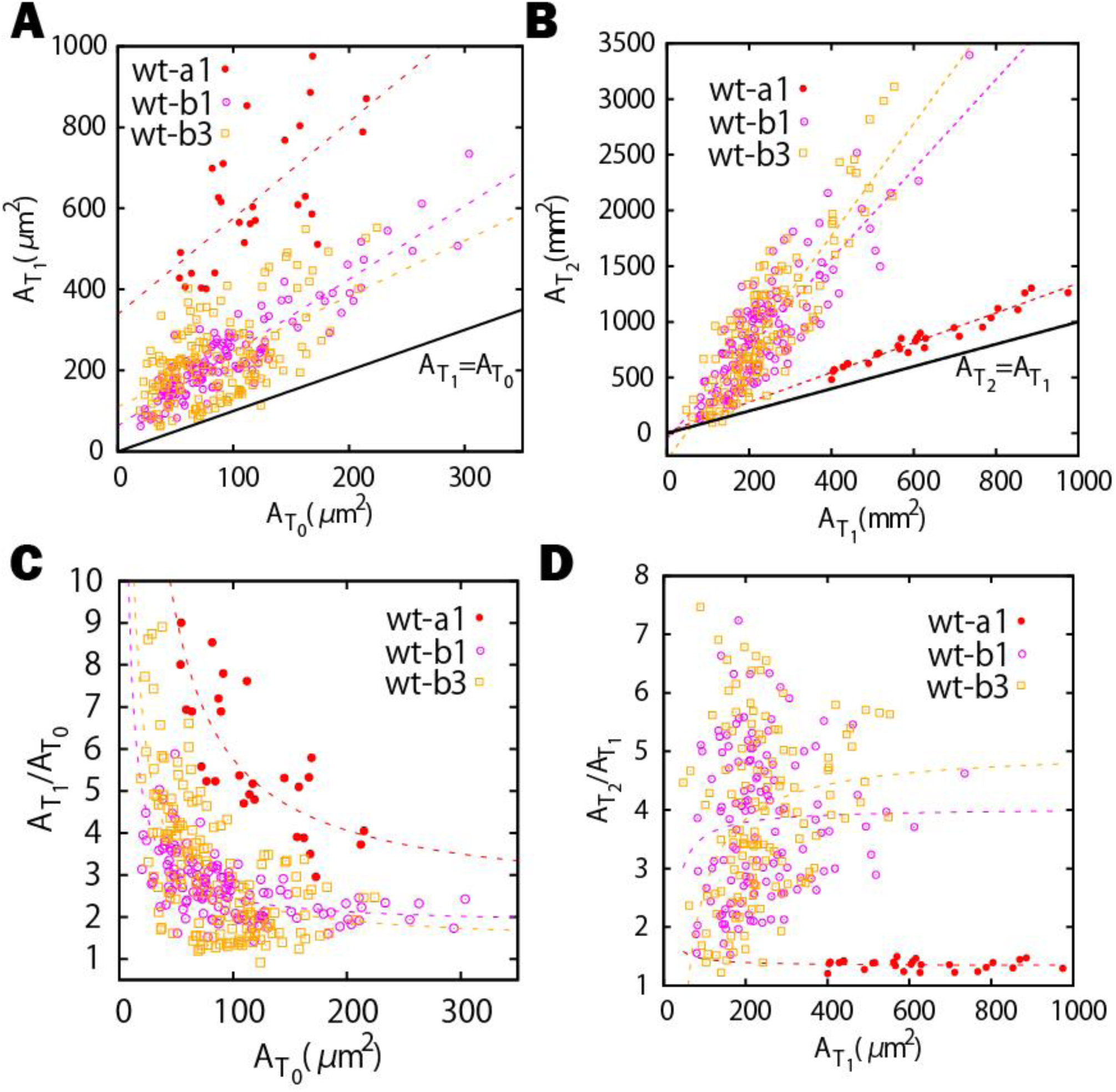
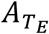 versus 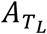 plot for the cell lineage groups at *T*_0_ versus those at *T*_1_ (A), and at *T*_1_ versus those at *T*_2_ (B). 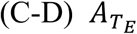 versus 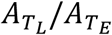 plot for the cell lineage groups at *T*_0_ versus those at *T*_1_ (C), and at *T*_1_ versus those at *T*_2_ (D). Error bars represent the 50%-confidence interval (see Materials and Methods).

**Fig. S3. (A-D-G-J).**
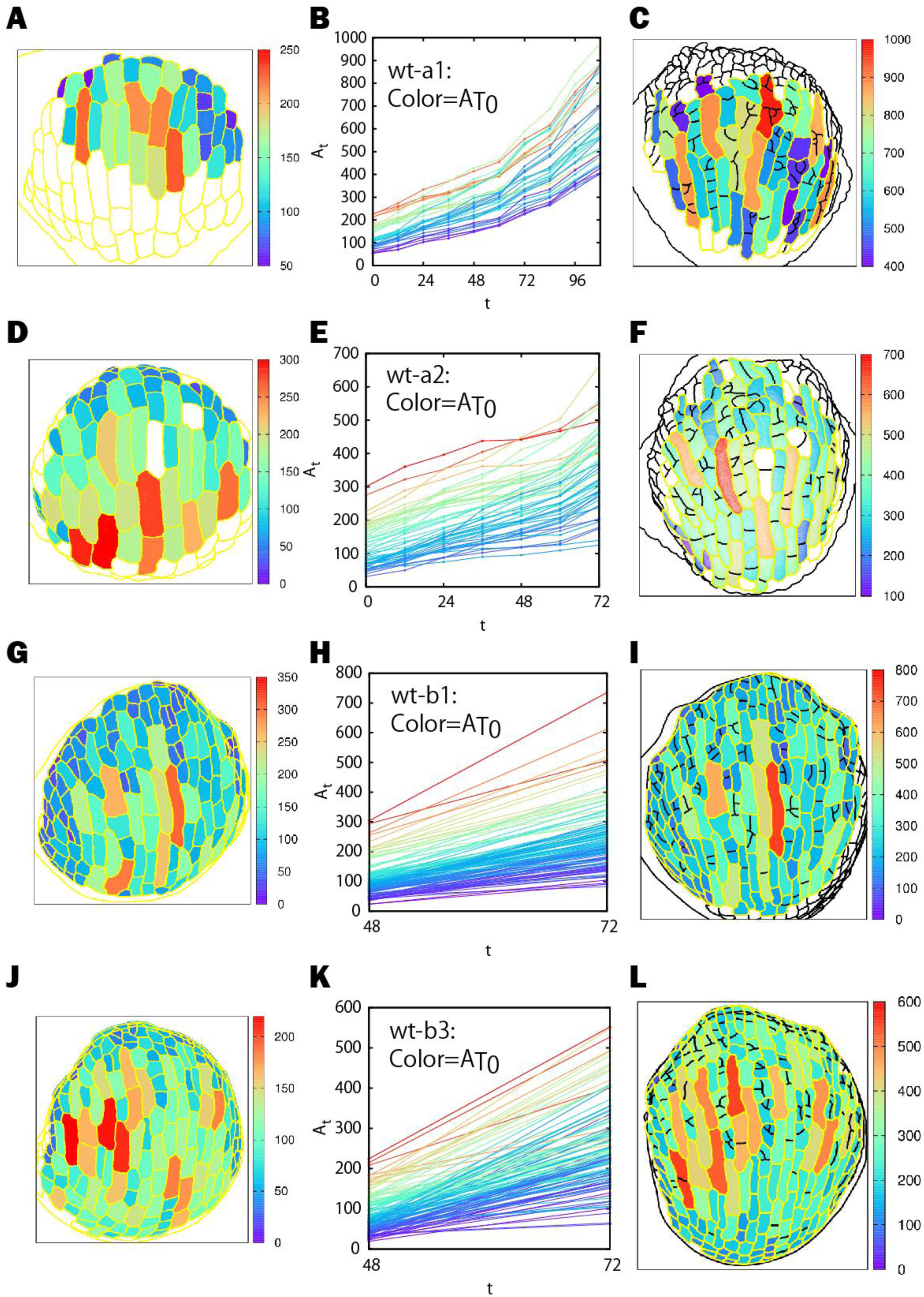
Heat map of the initial cell area 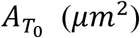. (B-E-H-K) Growth curves of cell lineage groups colored according to their initial size 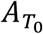. (C-F-I-L) Heat map of the size of the cell lineage group 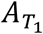 at a later time *T*_1_.

**Fig. S4. (A-D-G-J).**
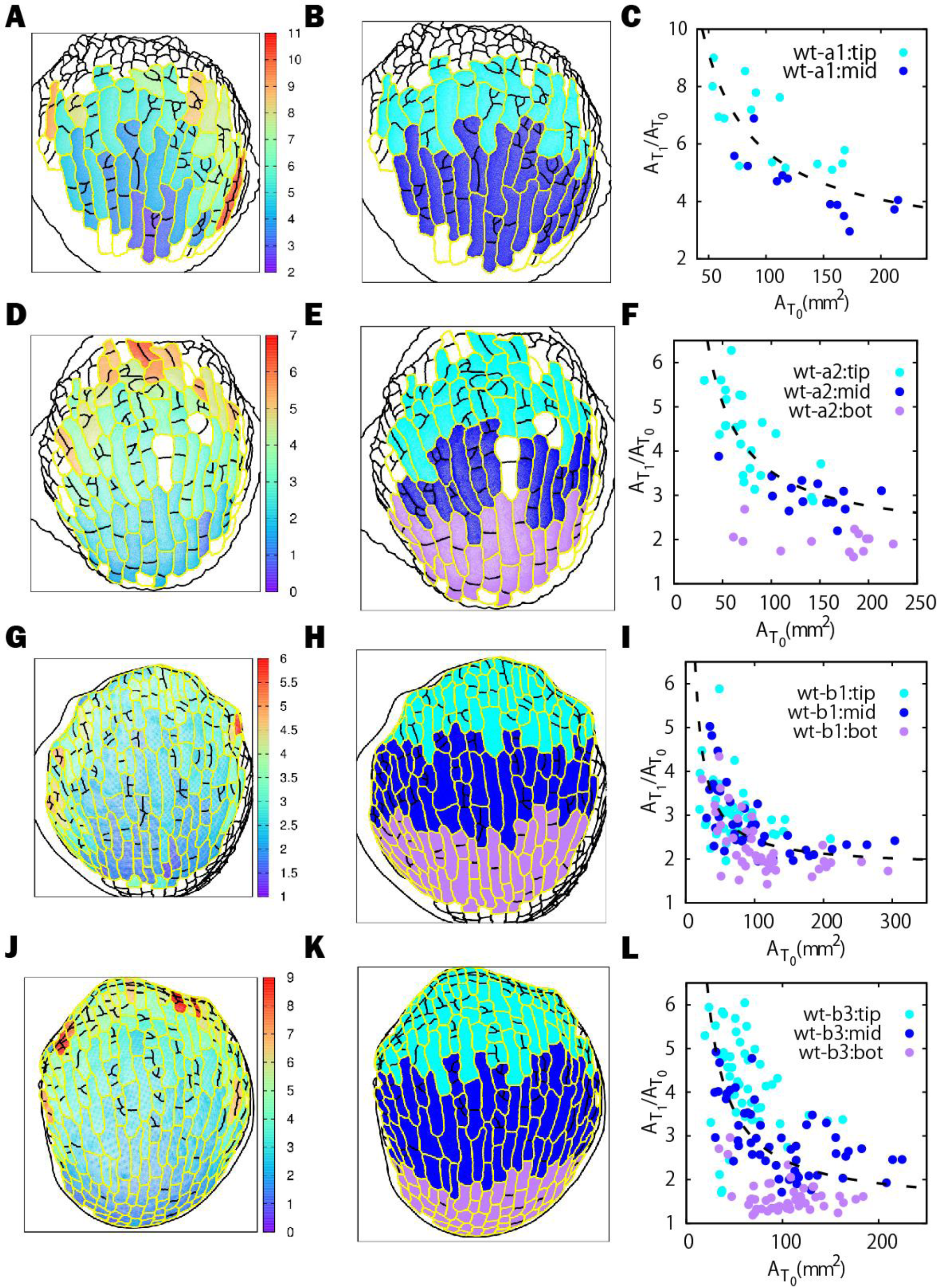
Heat map of the cumulative growth ratio 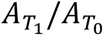 showing a continuous spatial trend from tip to bottom. (B-E-H-K) Classification of tip (cyan), middle (blue) and bottom (purple) regions. (C-F-I-L) Plot of cumulative growth ratios 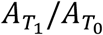 versus initial area 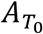 for cell lineage groups in each region.

**Fig. S5. (A-E).**
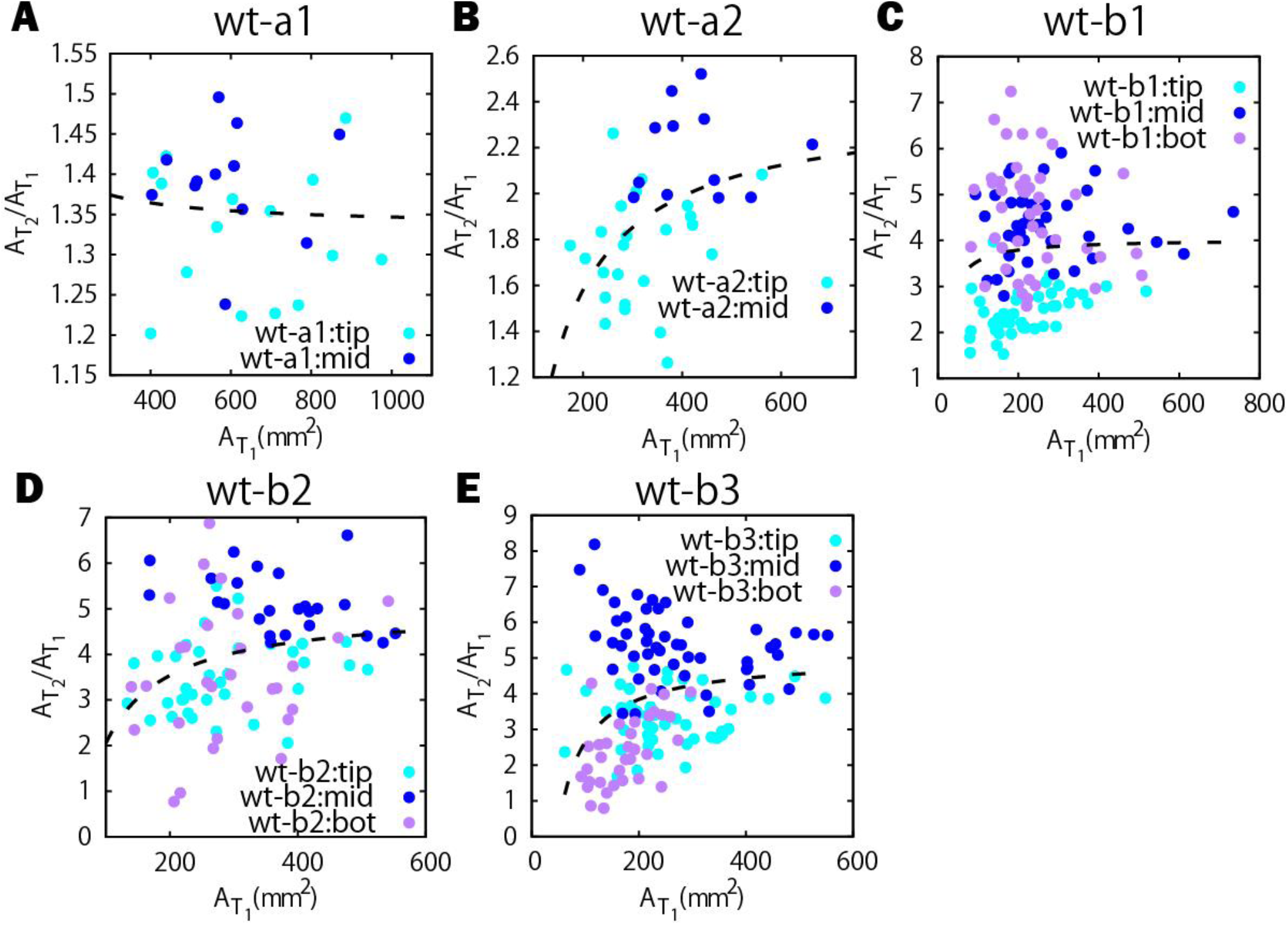
Plot of cumulative growth ratios 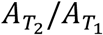 versus initial area 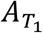 for cell lineage groups in each region for wt-a1, wt-a2, wt-b1, wt-b2 and wt-b3.

**Fig. S6. (A-B-C-D).**
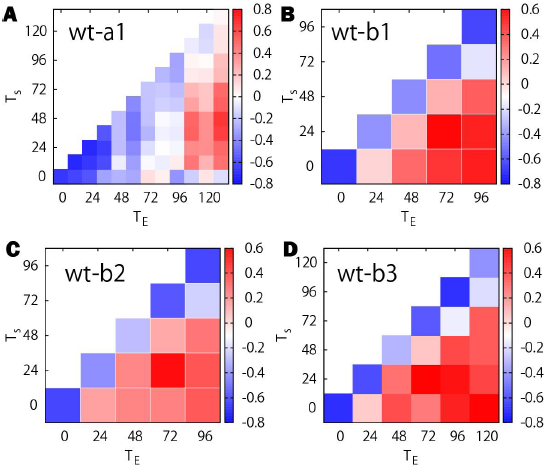
Spearman correlation coefficient between size and growth ratio for different *T*_*s*_ and *T*_*E*_ for the sepal wt-a1, wt-b1, wt-b2 and wt-b3, respectively.

**Fig. S7. (A-B-C-D).**
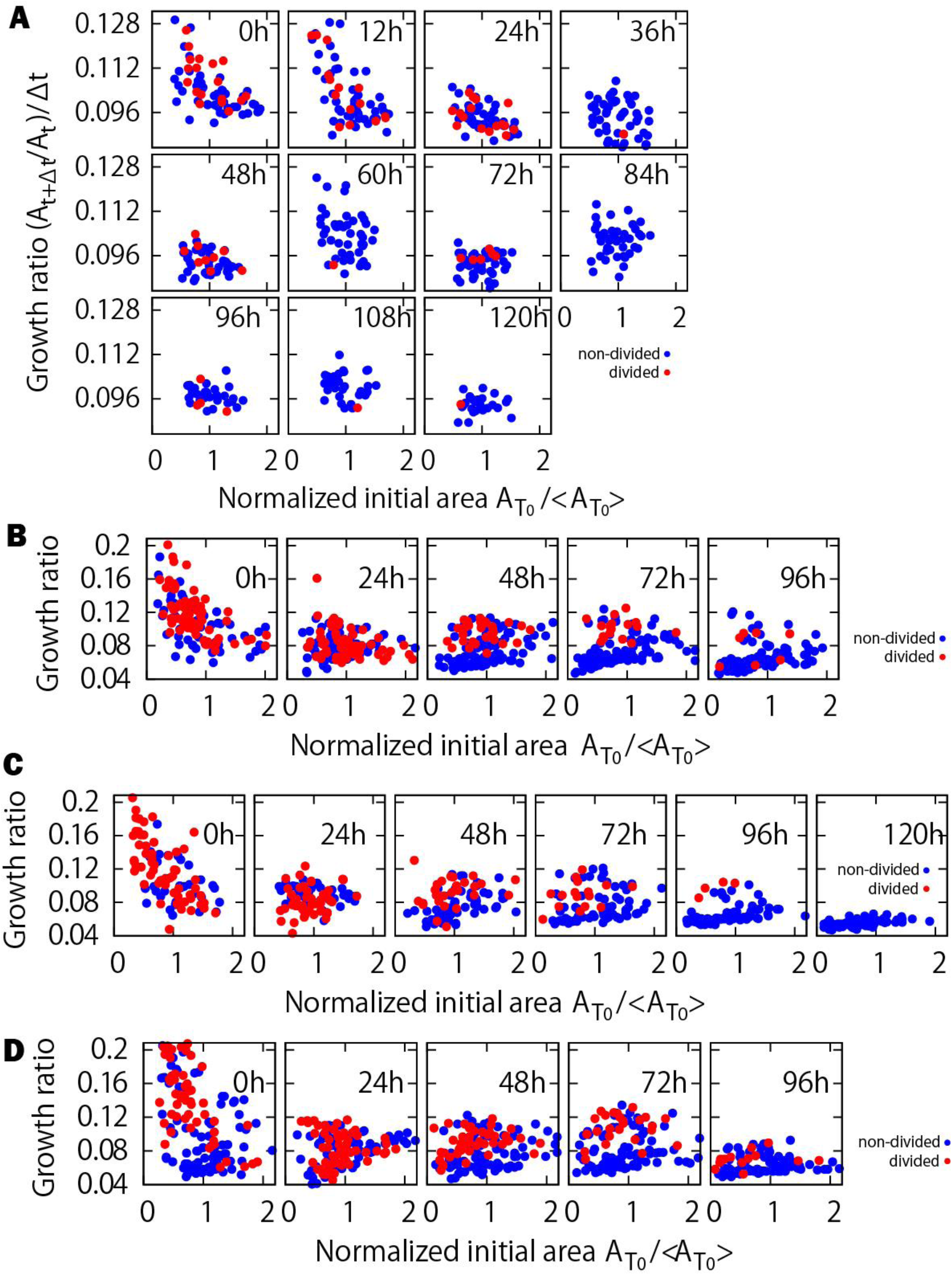
Plot of growth ratio at each time step (*A*_*t*+Δ*t*_/*A*_*t*_)/*Δt* versus normalized initial area 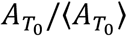 for the sepal wt-a1, wt-b1, wt-b2 and wt-b3, respectively.

**Fig. S8. (A-B-C-D).**
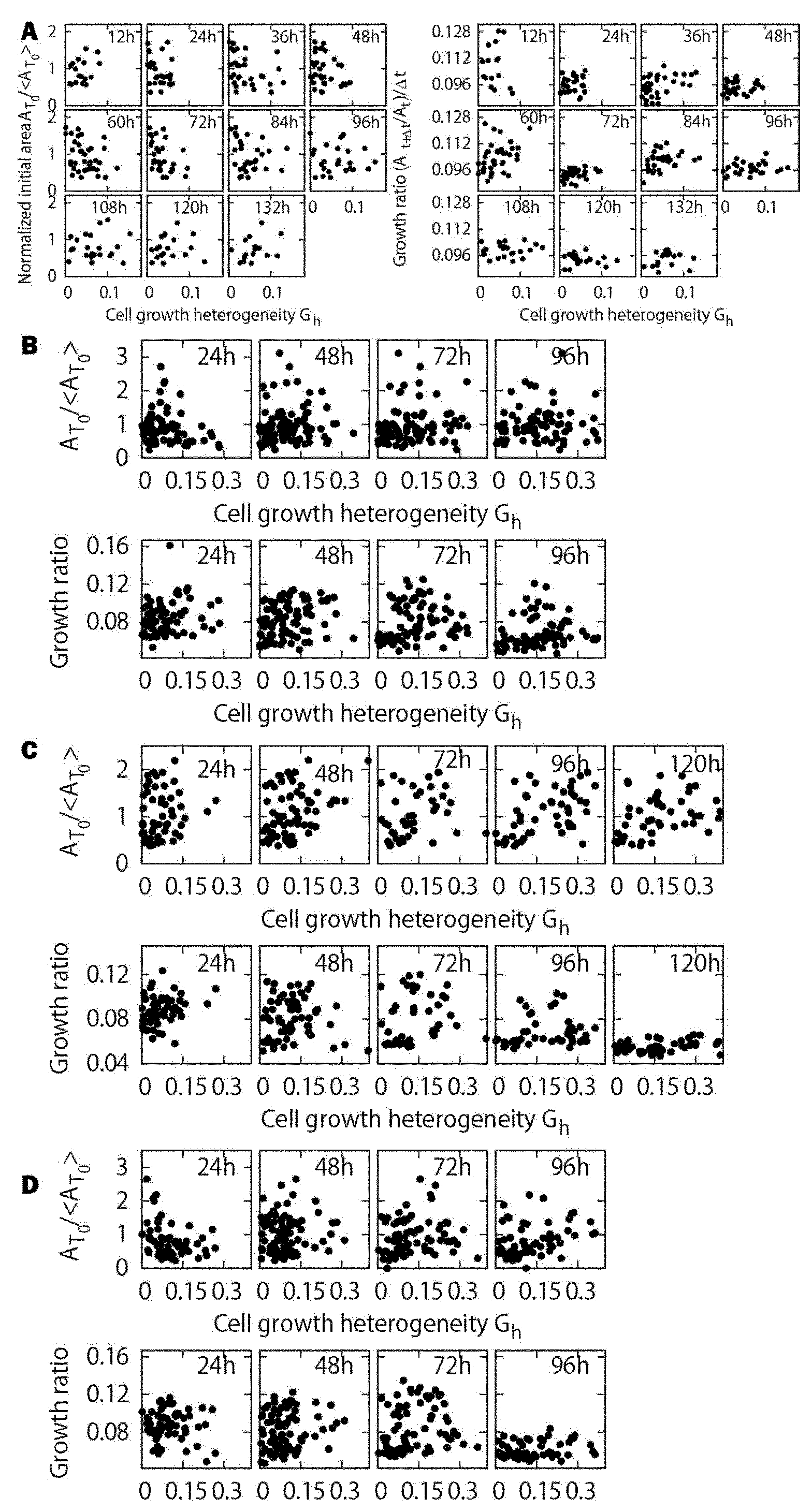
Cell growth heterogeneity versus normalized initial area and cell growth heterogeneity versus growth ratio of the cell lineage group at each time step for the sepal wt-a1, wt-b1, wt-b2 and wt-b3, respectively.

## References

1. Asl, L. K., Dhondt, S., Boudolf, V., Beemster, G. T. S., Beeckman, T., Inzé, D., Govaerts, W. and De Veylder L. (2011). Model-Based Analysis of Arabidopsis Leaf Epidermal Cells Reveals Distinct Division and Expansion Patterns for Pavement and Guard Cells. Plant Physiol. 156, 2172–2183.

2. Besson, S. and Dumais, J. (2011). Universal rule for the symmetric division of plant cells. Proc. Natl. Acad. Sci. 108(15), 6294–6299.

3. Cooke, J. (1975). Control of somite number during morphogenesis of a vertebrate, *Xenopus laevis*. Nature. 254, 196–199.

4. Barbier de Reuille, P., Routier-Kierzkowska, A-L., Kierzkowski, D., Bassel, G. W., Schüpbach, T., Tauriello, G., Bajpai, N., Strauss, S., Weber, A., Kiss, A., Burian, A., Hofhuis, H., Sapala, A., Lipowczan, M., Heimlicher, M. B., Robinson, S., Bayer, E. M., Basler, K., Koumoutsakos, P., Roeder, A. H. K., Aegerter-Wilmsen, T., Nakayama, N., Tsiantis, M., Hay, A., Kwiatkowska, D., Xenaios, L., Kuhlemeier, C. and Smith, R. S. (2015). MorphoGraphX: A platform for quantifying morphogenesis in 4D. eLife. 4, e05864.

5. Elsner, J., Michalski, M. and Kwiatkowska, D. (2012). Spatiotemporal variation of leaf epidermal cell growth: a quantitative analysis of *Arabidopsis thaliana* wild-type and triple cyclinD3 mutant plants. Ann. Bot. 109, 897–910.

6. Hervieux, N., Dumond, M., Sapala, A., Routier-Kierzkowska, A-L., Kierzkowski, D., Roeder, A. H. K., Smith, R. S., Boudaoud, A. and Hamant, O. (2016). A Mechanical Feedback Restricts Sepal Growth and Shape in *Arabidopsis*. Curr. Biol. 26, 1019–1028.

7. Hong, L., Dumond, M., Tsugawa, S., Sapala, A., Routier-Kierzkowska, A-L., Zhou, Y., Chen, C., Kiss, A., Zhu, M., Hamant, O., Smith, R. S., Komatsuzaki, T., Li, C-B., Boudaoud, and Roeder, A. H. K. (2016). Variable cell growth yields reproducible organ development through spatiotemporal averaging. Dev. Cell. 38(1), 15–32.

8. Kaplan, D. R. and Hagemann, W. (1991). The Relationship of Cell and Organism in Vascular Plants. BioScience, 41, 693–703.

9. Kierzkowski, D., Nakayama, N., Routier-Kierzkowska, A-L., Weber, A., Bayer, E., Schorderet, M., Reinhardt, D., Kuhlemeier, C. and Smith, R. S. (2012). Elastic Domains Regulate Growth and Organogenesis in the Plant Shoot Apical Meristem. Science. 335, 1096.

10. Marshall, W. F., Young, K. D., Swaffer, M., Wood, M., Nurse, P., Kimura, A., Frankel, J., Wallingford, J., Walbot, V., Qu, X. and Roeder, A. H. K. (2012). What determines cell size? BMC Biology, 10, 101.

11. Meyer, H. M. and Roeder, A. H. K. (2014). Stochasticity in plant cellular growth and patterning. Plant Evol. Dev. 5, 420.

12. Roeder, A. H. K., Chickarmane, V., Cunha, A., Obara, B., Manjunath, B. S. and Meyerowitz, E. M. (2010). Variability in the Control of Cell Division Underlies Sepal Epidermal Patterning in *Arabidopsis thaliana*. PLoS Biology. 8(5), e1000367.

13. Roeder, A. H. K., Cunha, A., Ohno, C. K. and Meyerowitz, E. M. (2012). Cell cycle regulates cell type in the *Arabidopsis* sepal. Development. 139, 4416–4427.

14. Smyth, D. R., Bowman, J. L. and Meyerowitz, E. M. (1990). Early Flower Development in Arabidopiss. The Plant Cell. 2, 7550767.

15. Spemann, H. and Mangold, H. (1924). Über Induktion von Embryonalanlagen durch, Implantation artfremder Organisatoren. Arch. mikrosk. Anat. EntwMech. 100, 599–638.

16. Tauriello, G., Meyer, H. M., Smith, R. S., Koumoutsakos, P. and Roeder, A. H. K. (2015). Variability and constancy in cellular growth of Arabidopsis sepals. Plant Physiol. 169(4), 2342–2358.

17. Tsukaya, H. 2003. Organ shape and size: a lesson from studies of leaf morphogenesis. Curr. Opin. Plant Biol. 6, 57–62.

18. Uyttewaal, M., Burian, A., Alim, K., Landerin, B., Borowska-Wykręt, D., Dedieu, A., Peaucelle, A., Ludynia, M., Traas, J., Boudaoud, A., Kwiatkowska, D. and Hamant, O. (2012). Mechanical Stress Acts via Katanin to Amplify Differences in Growth Rate between Adjacent Cells in *Arabidopsis*. Cell. 149, 439–451.

19. Willis, L., Rafahi, Y., Wightman, R., Landrein, B., Teles, J., Huang, K. C., Meyerowitz, E.M. and Jönsson, H. (2016). Cell size and growth regulation in the *Arabidopsis thaliana* apical stem cell niche. PNAS, 113, 8238–8246.

20. Yang, W., Schuster, C., Beahan, C. T., Charoensawan, V., Peaucelle, A., Bacic, A., Doblin, M. S., Wightman, R. and Meyerowitz, E. M. (2016). Regulation of meristem morphogenesis by cell wall synthases in Arabidopsis. Current Biology, 26(11), 1404–1415.

